# Functional Impact of Glycosylation and Splicing Variants of LRIG1 on EGFR Proteostasis in Cancer

**DOI:** 10.1101/2025.02.05.636756

**Authors:** Jumpei Kondo, Himari Nagao, Koki Oyama, Daisuke Sakon, Honoka Nakayama, Shinji Takamatsu, Eiji Miyoshi

## Abstract

LRIG1, a membrane glycoprotein, has emerged as a significant stem cell marker and negative regulator of receptor tyrosine kinases (RTKs), including EGFR. Glycosylation is a major post-translational modification, which plays a crucial role in protein function and stability. In cancer biology, abnormal glycosylation can serve as a biomarker and contribute to pathogenesis. We aimed to investigate the effects of glycosylation on LRIG1 functions. Through database analysis and experimental approaches, we focused on evolutionarily conserved glycosylation sites of LRIG1, particularly N74 in humans. We found that a mutation of the N74 glycosylation site (N74Q) enhances LRIG1’s binding to EGFR and promotes EGFR degradation. Furthermore, we identified a naturally occurring splice variant of LRIG1 lacking exon 2, which includes the N74 site, that shows similar enhanced EGFR binding and degradation. Analysis of TCGA data revealed that exon 2 skipping in LRIG1 occurs in various cancers, with higher levels correlating with poorer survival in breast and ovarian cancers. Our findings suggest that the absence of glycosylation at N74 enhances LRIG1-EGFR binding, providing an example of glycosylation negatively regulating protein-protein interaction. This mechanism, achieved through alternative splicing, provides insights into the importance of glycosylation deficiency in cancer biology.

## Introduction

The membrane glycoprotein LRIG1 has recently gained attention as a stem cell marker in various tissues such as intestinal epithelium^1^, epidermis^2^, corneal^3^, and neurons^4^. Functionally, LRIG1 is recognized as a negative regulator of receptor tyrosine kinases (RTKs), including EGFR and other ERBB family receptors^1,5^, as well as MET^6^. Mechanistically, LRIG1 interacts with these RTKs on the cell membrane, and when a ligand binds to the RTK, LRIG1 recruits the E3 ubiquitin ligase Cbl, which induces lysosomal degradation of the target receptor through ubiquitination^7^. Through its role in regulating receptor tyrosine kinases (RTKs), LRIG1 contributes to differentiation control and the maintenance of stem cell quiescence^4,8^. In cancer contexts, high levels of LRIG1 are associated with a favorable prognosis in lung and prostate cancers, where it plays a role in suppressing tumor growth^9^. Conversely, during developmental stages characterized by vigorous cell proliferation, LRIG1 can control the direction of differentiation without suppressing cell proliferation^10^. Notably, high LRIG1 expression is also observed in rapidly proliferating intestinal epithelial transit-amplifying (TA) cells^1,5^. These observations suggest that the degradation effect of LRIG1 on RTKs is regulated by both its expression level and the activity of its degradation-inducing mechanisms, although specific details of the latter regulation are still unclear.

Glycans, often referred to as carbohydrates or sugar chains, are complex biomolecules that play pivotal roles in numerous biological processes^11,12^. These structures are covalently attached to proteins and lipids in a process known as glycosylation, a post-translational modification that occurs in nearly all eukaryotic cells. Glycosylation is highly diverse, producing a vast array of glycan structures that contribute to biomolecular complexity beyond the genome or proteome. Glycan chains of glycoproteins include O- and N-glycoside-linked glycans; O-glycans are attached to serine and threonine, while N-glycans are linked to asparagine^11–13^. The addition of N-glycosidic linkages, termed N-glycosylation, is also essential for the proper protein folding and localized trafficking of membrane and secretory proteins^14,15^.

At the cellular community level, glycans function as vital mediators of cellular communication, structural integrity, and molecular recognition^15^. For instance, cell surface glycans play a pivotal role in dictating cell-cell interactions^14,16^, immune recognition^17^, and pathogen adhesion^18,19^. Aberrant glycosylation is frequently observed in diseases such as cancer, autoimmune disorders, and neurodegenerative conditions, underscoring its critical role in pathology^11,14,15^. A specific example is the epidermal growth factor receptor (EGFR), which promotes proliferation in various cancers, including lung and colon cancer. Its function is modulated by the addition of fucose to glycans, a process known as fucosylation^20^. Despite the fundamental role of glycosylation in protein function, its contribution to LRIG1 remains largely unexplored, with only a single report suggesting its potential importance in cellular quiescence^5^. This raises the possibility that regulation of LRIG1 glycosylation could be a factor influencing its functions; however, there are no other reports on this topic.

In the present study, we investigated the effects of LRIG1 glycosylation on its ability to bind to EGFR and induce degradation by analyzing databases and examining LRIG1 glycosylation site mutants. Additionally, we explored the potential for changes in glycosylation sites in actual cancer cases and whether these changes may relate to prognosis.

## Methods

### Cell lines and organoids

Human embryonic kidney cell line HEK293T and human colon adenocarcinoma cell line Caco-2 were purchased from American Type Culture Collection (ATCC, Manassas, VA). HEK293T / FT cells were cultured in DMEM High Glucose (Dulbecco’s Modified Eagle’s medium; nacalai tesque, Kyoto, Japan) supplemented with 10% FBS (Fetal bovine serum; NICHIREI BIOSCIENCES INC, Tokyo, Japan) and 100 U/mL penicillin, 100 μg/mL streptomycin (nacalai tesque). Caco-2 cells were cultured in DMEM (nacalai tesque) supplemented with 20% FBS and 100 U/mL penicillin, 100 μg/mL streptomycin. These cells were cultured under humidified conditions at 37°C with 5% CO₂ (Astec, Fukuoka, Japan; CPI-165).

Human colorectal cancer organoid C45 derived from a clinical specimen^21^ was provided by the Department of Clinical Bio-resource Research and Development, Graduate School of Medicine, Kyoto University (Dr. Masahiro Inoue). The organoids were cultured in a 1:7 mixture of StemPro hESC SFM (Gibco, Palo Alto, CA) and Advanced DMEM/F12 (Gibco) supplemented with L-Alanyl-L-Glutamine Solution (Wako, Osaka, Japan). Organoid-based experiments were carried out with the approval of the Ethics Committee of Osaka University Hospital.

### Vector construction and transfection

The coding sequence (cds) of LRIG1 was cloned from Caco-2 cells and C45 organoids. cDNA was synthesized using the PrimeScriptTM II 1st strand cDNA Synthesis Kit (Takara Bio, Kusatsu, Shiga) from mRNA extracted with the RNeasy® Mini Kit (QIAGEN, Hilden, Germany). The PCR products amplified using PrimeSTAR Max DNA Polymerase (Takara Bio) and pCI-neo (Promega, Madison, WI) were treated with restriction enzymes EcoRI and SalI (Takara Bio). After purification using NucleoSpin Gel and PCR Clean-up (MACHEREY-NAGEL GmbH & Co., Duren, Germany), ligation was carried out with DNA Ligation Kit <Mighty Mix> (Takara Bio), and vectors pCI-neo-LRIG1 and C45-pCI-neo-LRIG1, derived from Caco-2 and C45 respectively, were obtained.

For the C-terminal flag tagging of LRIG1 in pCI-neo-LRIG1, a modified Gibson assembly method using NEBuilder (New England Biolabs, Beverly, MA) was employed. Fragments were amplified using PrimeSTAR Max DNA Polymerase. To generate LRIG1 variants lacking N-glycosylation sites, PCR-based mutagenesis was performed using the pCI-neo-LRIG1-Flag vector as a template with PrimeSTAR Max DNA Polymerase. LRIG1 sequences in all the generated vectors were confirmed by Sanger sequencing using Big-dye terminator v3.1 (Thermo Fischer Scientific, Waltham, MA). Primers used for vector construction were listed in Supplementary Table S1. pHAGE-EGFR was a gift from Gordon Mills & Kenneth Scott (Addgene plasmid # 116731).

For transient gene expression, transfection was performed on adherent cells seeded in a 6- well plate using a mixture of 1 µg plasmid, 6 µL X-tremeGENE HP DNA Transfection Reagent (Roche, Basel, Switzerland), and 100 µL Opti-MEM (Thermo Fisher Scientific). Cells 48 h after transfection were used for experiments.

### Western blotting

Cell pellets were lysed in TNE buffer (10 mM Tris-HCl pH 7.8, 1% NP40, 0.5 M NaCl, 1 mM EDTA) supplemented with PIC (Protease Inhibitor Cocktail; nacalai tesque). After sonication and centrifugation, the supernatant was collected as whole-cell lysate. For the deglycosylation assay, lysates were treated additionally by PNGase F (New England Biolabs, Ipswich, MA) under denaturing conditions according to the manufacturer’s protocol. The lysate was mixed with 4× SDS sample buffer containing 2-mercaptoethanol (nacalai tesque), boiled, and subjected to SDS-PAGE. Subsequently, proteins were transferred to Polyvinylidene fluoride (PVDF; Millipore) membranes for western blotting. The primary and secondary antibodies used are listed in Supplementary Table S2. For band detection, Chemi-Lumi One Super (Nacalai) was used, and chemiluminescence signals were detected using the FUSION Chemiluminescence imaging system (Vilber-Lourmat, Collegien, France).

### Immunoprecipitation

For evaluation of LRIG1-EGFR binding, whole cell lysates were diluted in TBS (2.5 mM Tris, 13.8 mM NaCl, 0.27 mM KCl) to a protein concentration of 400 µg/500 µL. For the detection of EGFR ubiquitination, cells were lysed using SDS-free RIPA buffer (50 mM Tris-HCl pH 7.4, 1% NP40, 150 mM NaCl, 0.5% Sodium deoxycholate, 1 mM EDTA) supplemented with PIC and 10 mM NEM (N-Ethylmaleimide; nacalai tesque), then diluted in TBS at 400 µg/500 µL. Pre-clearing was performed using control mouse IgG (PeproTech, Cranbury, New Jersey) and SureBeads Protein G Magnetic beads (BIO-RAD). Subsequently, the samples were incubated with anti-Flag (Sigma-Aldrich) or anti-EGFR antibody (CST) at room temperature for 1 hour, followed by capture with magnetic beads. The immunoprecipitates were then eluted by adding 4× SDS sample buffer containing 2-mercaptoethanol and boiling. The eluted samples were evaluated by western blotting as described above.

### Database Analysis

The ortholog sequences of LRIG1 in vertebrates were searched using the NCBI database (https://www.ncbi.nlm.nih.gov/), and alignments were performed using the COBALT: Multiple Alignment Tool on the same database. The selection of 22 representative species from 635 vertebrate species registered in the database was carried out prioritizing species with high-quality genome information, covering major taxonomic groups, and including species at evolutionary significant positions.

For the analysis of glycosylation sites, searches were conducted using four databases linked to different mass spectrometry data: Glygen (https://glygen.org/), GlycoProtDB (https://acgg.asia/db/gpdb/), PeptideAtlas (https://peptideatlas.org/), and N-GlycositeAtlas (http://nglycositeatlas.biomarkercenter.org/). Information on human and mouse LRIG1 was collected from the glycoproteomics analysis data registered in each database, and it was determined whether peptides containing each glycosylation site were detected.

TCGA SpliceSeq (https://bioinformatics.mdanderson.org/TCGASpliceSeq/)^22^ was used for the analysis of exon skipping based on RNA-seq data registered in TCGA (The Cancer Genome Atlas Program). The exon 2 skipping event of LRIG1 is registered as LRIG_ES_192431. In the SpliceSeq algorithm, PSI (Percent Spliced In) values are calculated when there are 8 or more reads related to Alternative Splicing. Analysis was conducted based on data obtained from the above database for six types of cancer where more than 10% of samples were assigned PSI values for LRIG_ES_192431. Disease-specific and progression-free survival analysis was performed on OncoSplicing (http://www.oncosplicing.com/)^23^.

## Results

### Database analysis revealed evolutionarily conserved glycosylated sites on LRIG1

The human LRIG1 protein contains six N-glycosylation consensus sequences, Asn-X-Thr or Asn-X-Ser. Asparagines that undergo glycosylation tend to be more evolutionarily conserved compared to non-glycosylated asparagines^24^. In addition, functionally important glycosylation sites are conserved in certain glycoproteins^25–27^. Thus, to predict functionally significant N-glycosylation sites on LRIG1, we performed a cross-species analysis of conserved consensus sequences, focusing on 22 representative vertebrate species. In mammals, all six N-glycosylation sites were highly conserved (Fig. 1A). However, only the first, third, and sixth glycosylation sites from the N-terminus (corresponding to N74, N246, and N684 in humans) were found to be conserved among vertebrates (Fig. 1A). For the fifth glycosylation sequence (corresponding to N318 in humans), no glycosylation sequence was observed at the homologous position in the High Himalaya frog, Elephant shark, and Whitespotted bamboo shark. Nevertheless, a glycosylation sequence was found five amino acids upstream in these species, suggesting that glycosylation near this site may also play a functional role. To verify the glycosylation of these sites, we re-evaluated mass spectrometry data from publicly available databases, including Glygen, GlycoProtDB, PeptideAtlas, and N-GlycositeAtlas. Our analysis revealed glycan modifications at the first, second, and fifth glycosylation sites from the N-terminus of LRIG1 (N74, N150, N318 in humans) (Table 1). Based on these findings, we hypothesized that glycan modifications in the extracellular domain of LRIG1 protein—particularly at the first and the fifth glycosylation sites (N74 and N318 in humans), which were corroborated by two database analyses—hold significant functional importance. Consequently, we conducted the following experiments to test this hypothesis.

**Figure 1.**
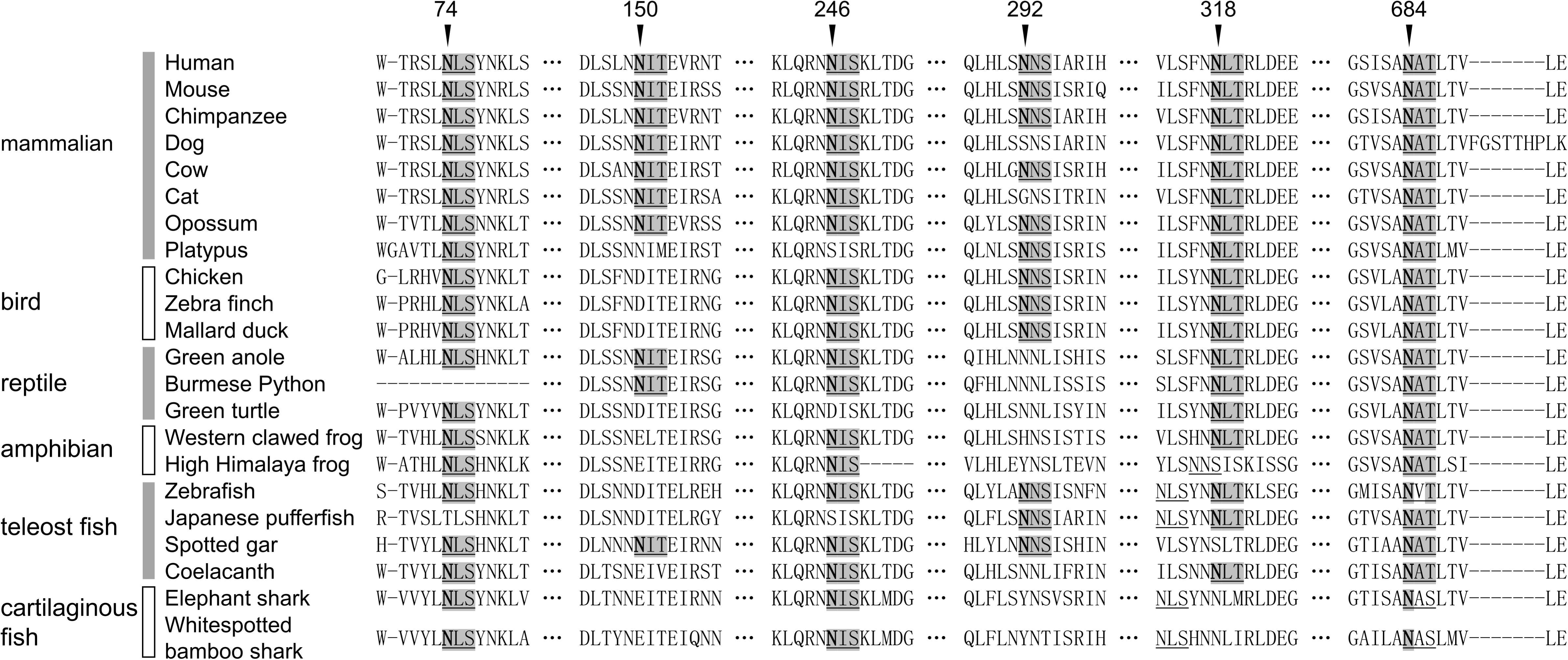
Evolutionarily conserved glycosylation sites on LRIG1. The evolutionary conservation of N-glycosylation sites on LRIG1 was investigated through the analysis of amino acid sequences of LRIG1 orthologs from a range of representative vertebrate species. The classes and species are displayed on the left. The numbers represent the amino acid positions in humans, and the use of underlines indicates the presence of glycosylation sequences. Hyphens: gaps in the data, horizontal ellipsis: omission between sites, shaded letterings: glycosylation sequences conserved in humans, bold letters: asparagine residues to which glycans can be attached.

**Table 1.**
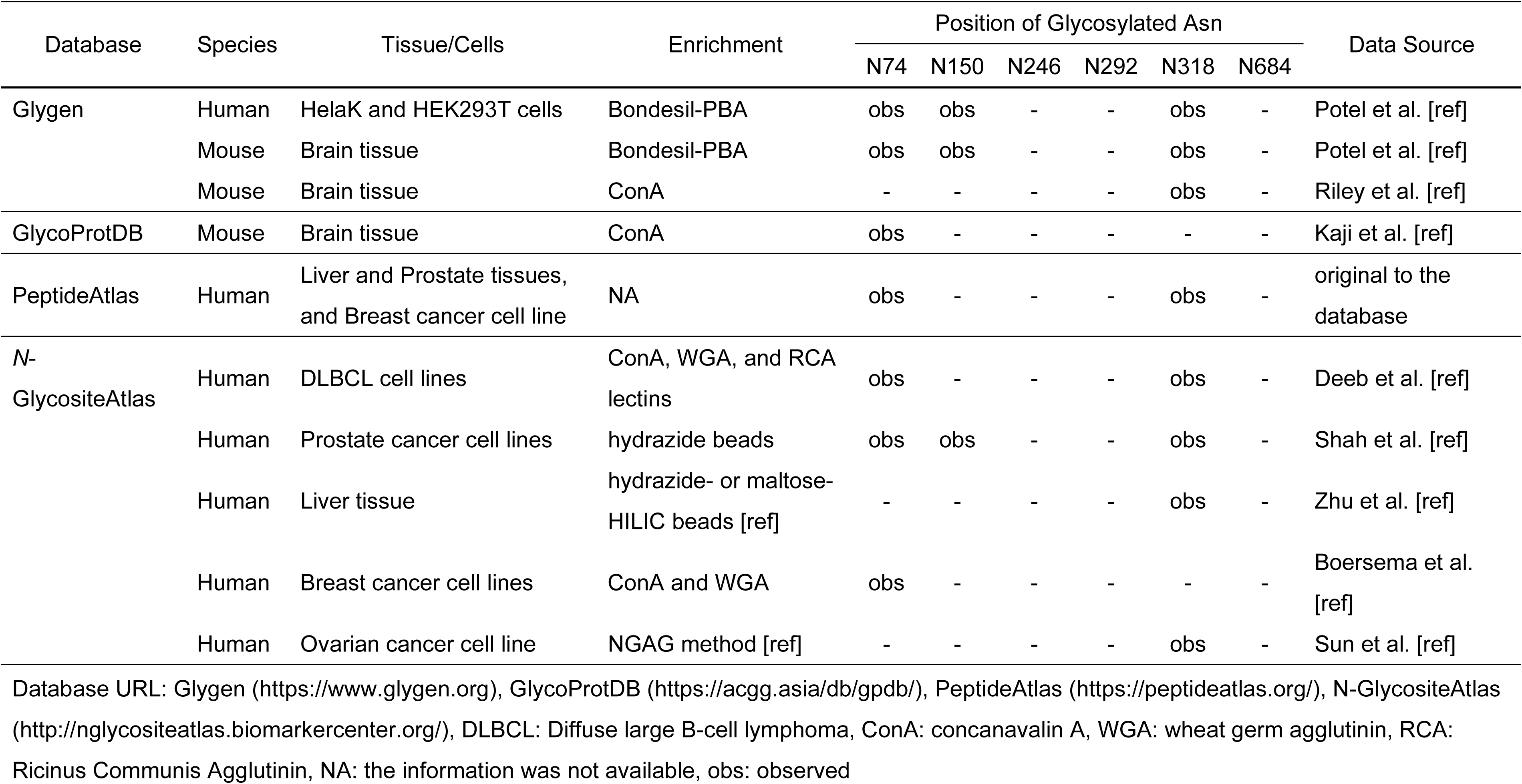
Putative glycosylation sites with actual glycosylation that were detected by glycoproteomics

### Glycosylation of LRIG1 affects interaction with EGFR and its degradation

Firstly, we confirmed that LRIG1 expressed in HEK293T cells is glycosylated. Lysates were treated with PNGase F, an enzyme that removes the entire glycan from glycoproteins. Western blot analysis showed a shift to a lower molecular weight in LRIG1, as well as in EGFR, a well-characterized glycoprotein used as a positive control (Fig. 2A). Next, to investigate the functional significance of each glycosylation site, Asn-to-Gln (N>Q) mutations were introduced into LRIG1 cloned from human cells, generating glycosylation-deficient mutants. LRIG1 cDNA was cloned from CACO2 cells, which were confirmed via COSMIC to have no mutations other than single nucleotide variants (SNVs). The coding sequence (cds) of the obtained clone was verified by Sanger sequencing to contain only non-pathogenic SNVs registered in dbSNP (https://www.ncbi.nlm.nih.gov/snp/). However, the N292Q mutant could not be generated due to limitations in primer design for PCR-based mutagenesis. Consequently, we focused on the remaining five mutants (Fig. 2B).

**Figure 2.**
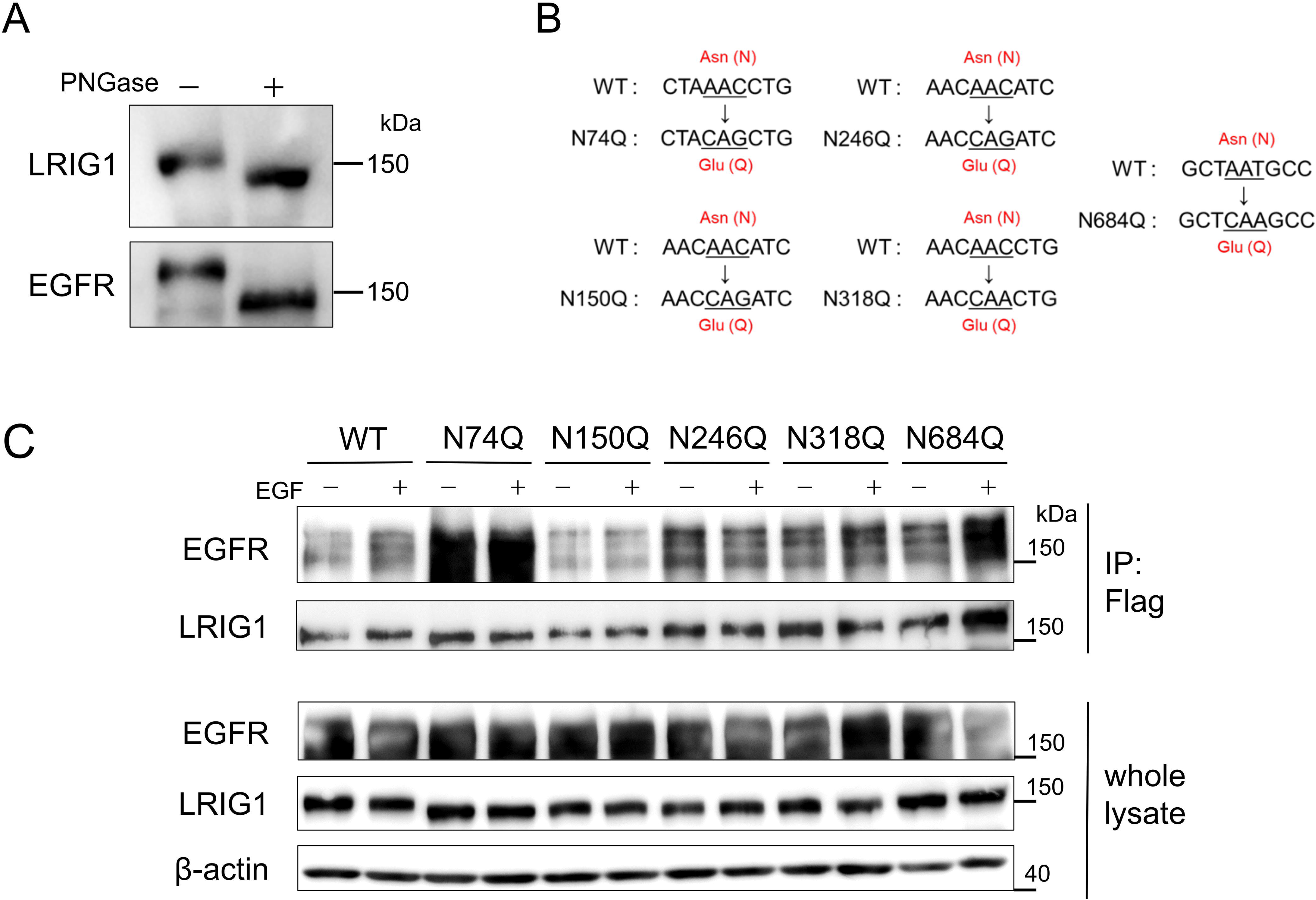
Glycosylation sites of LRIG1 regulate the binding with EGFR. A. Immunoblotting analysis of HEK293T cells with LRIG1 overexpression. Cell lysate in the right lane was treated with PNGase. Proteins were detected with antibodies for indicated proteins. B. Schemas of LRIG1 mutants in five glycosylation sites. C. Immunoblotting analysis of HEK293T cells. Cells were transfected with WT or mutant LRIG1-Flag and EGFR vectors and stimulated with 20 ng/mL recombinant human EGF. The upper two blots are for proteins co-immunoprecipitated with Flag antibody. The lower three blots are for whole lysates.

Upon overexpression of these mutants in HEK293T cells, Western blot analysis showed that all mutants exhibited bands with lower molecular weights compared to wild-type (WT) LRIG1. This effect was most apparent in the N74Q mutant, confirming that these sites are indeed glycosylated (Fig. 2C). Given that LRIG1-EGFR binding is an important step for LRIG1-mediated EGFR degradation, co-immunoprecipitation (co-IP) was performed using FLAG-tagged LRIG1 to quantify EGFR binding (Fig. 2C). The N74Q mutant exhibited enhanced EGFR binding, irrespective of EGF stimulation, while the N684Q mutant showed increased binding only upon EGF stimulation. In contrast, the N318Q mutant, which was likewise implicated as functionally relevant based on database analysis (Figure 1), did not show any alteration in its ability to bind EGFR.

To further examine the functional relevance of N74 glycosylation, we evaluated the ability of the N74Q mutant to promote EGFR degradation. Transient expression of WT or N74Q LRIG1 in HEK293T cells, followed by EGF stimulation, revealed that the N74Q mutant enhanced EGFR degradation compared to the WT (Figs. 3A, B). Given that EGF-induced EGFR degradation proceeds via lysosomal pathways mediated by ubiquitination, immunoprecipitation experiments showed that the N74Q mutant promoted EGF-stimulated EGFR ubiquitination (Fig. 3C). Together, these findings suggest that the absence of glycosylation at N74 enhances LRIG1-EGFR binding, thereby facilitating EGF-stimulated EGFR degradation.

**Figure 3.**
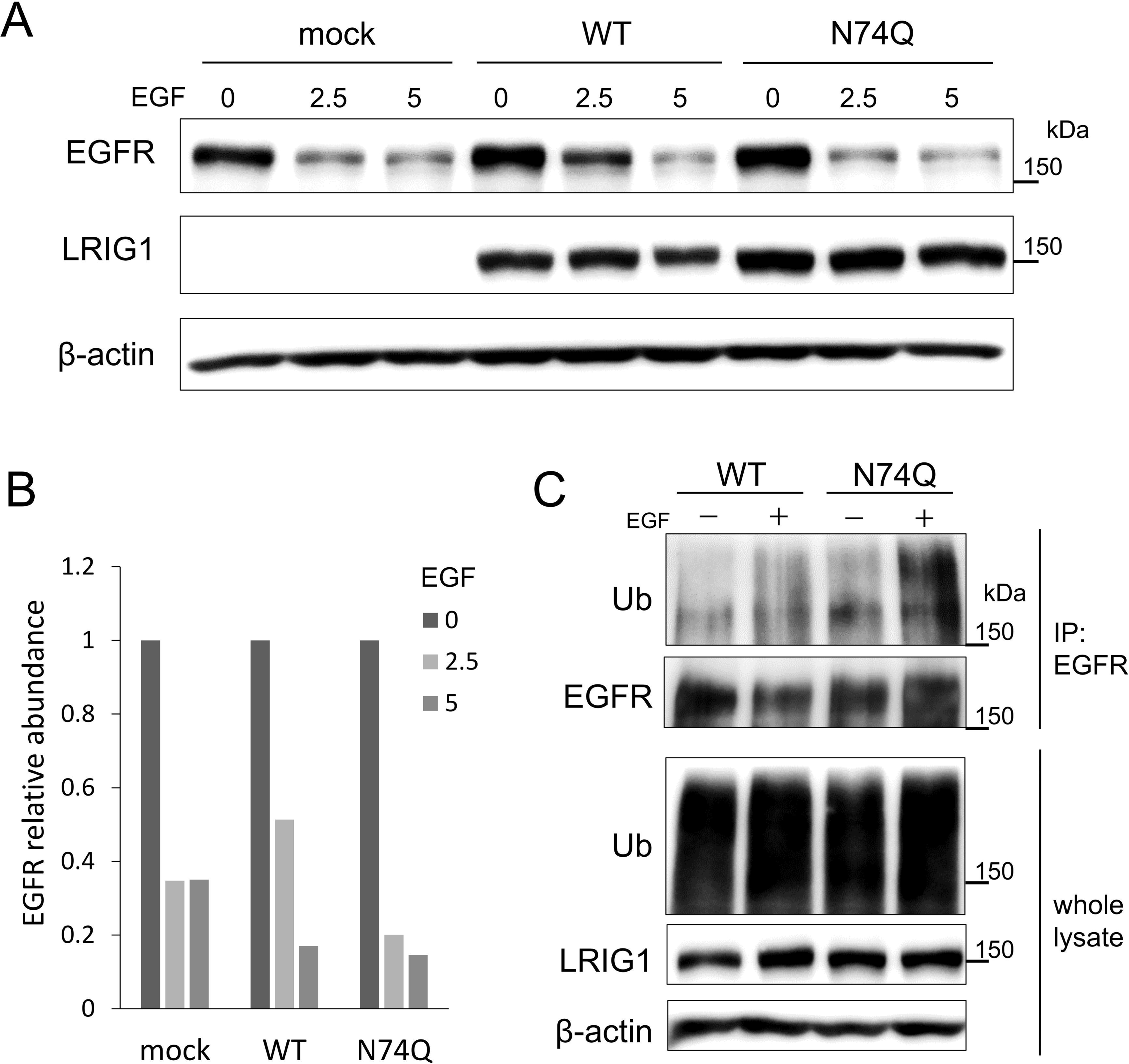
Glycosylation of LRIG1 at N74 regulates the degradation rate of EGFR. A. Immunoblotting analysis of HEK293T cells transfected with WT or N74Q mutant LRIG1-Flag vectors and stimulated with recombinant human EGF at indicated concentrations (ng/mL). B. Semi-quantification of the band in Figure 3A. C. Immunoblotting analysis of HEK293T cells. Cells were transfected with WT or N74Q mutant LRIG1-Flag and stimulated with 10 ng/mL recombinant human EGF. The upper two blots are for proteins immunoprecipitated with EGFR antibody. The lower three blots are for whole lysates.

### A splicing variant of LRIG1 lacking exon 2 promotes interaction with and degradation of EGFR

Alternative splicing often leads to exon skipping, which can alter protein function. Although examples involving the loss of N-glycosylation sites are rare, their functional significance has been noted in specific instances, such as Neuropilin-1 (NRP1)^28^. To investigate whether LRIG1 undergoes similar splicing events that disrupt glycosylation and affect its function, we explored the potential existence of naturally occurring LRIG1 splicing variants. In human LRIG1, the N74 glycosylation site is encoded in exon 2. In our experiment, one out of four LRIG1 coding sequences (cds) obtained from the colorectal cancer organoid C45 was identified as an exon 2 skipping variant (Δex2) (Fig. 4A, clone #14). To assess the functional impact of this splicing variant, we overexpressed full-length LRIG1 derived from Caco-2 and C45, as well as Δex2 LRIG1 derived from C45, in HEK293T cells. The ability of these variants to degrade EGFR following EGF stimulation was evaluated. The Δex2 LRIG1 variant exhibited enhanced EGFR degradation activity compared to the full-length LRIG1 (Fig. 4B, C). Furthermore, when co-expressed with EGFR in HEK293T cells, immunoprecipitation analysis revealed that Δex2 LRIG1 bound more strongly to EGFR than full-length LRIG1 (Fig. 4D). These findings indicate that Δex2 LRIG1, which lacks the N74 glycosylation site, has an enhanced EGFR-binding affinity and an increased capacity to induce EGFR degradation, reflecting the functional effects observed with the N74Q mutant of LRIG1.

**Figure 4.**
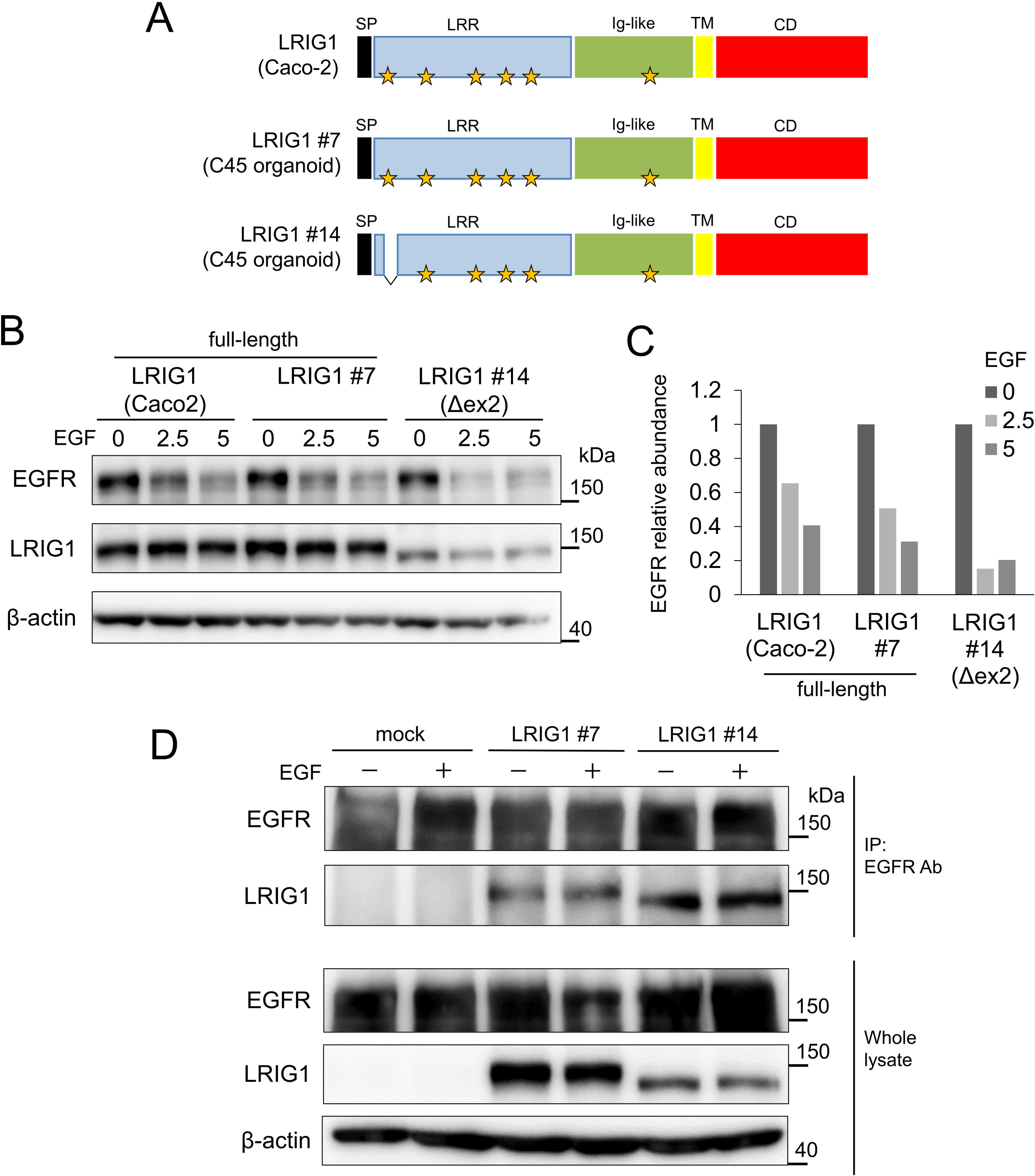
A splicing variant of LRIG1 lacking exon 2 promotes interaction with and degradation of EGFR. A. Schemas of LRIG1 clones derived from Caco-2 and C45 organoids. The regions on the cds corresponding to the LRIG1 protein domain are shown in different colors. Yellow stars indicate glycosylation sites. SP: signal peptides, LRR: leucine-rich repeats, Ig-like: immunoglobulin-like domains, TM: transmembrane domain, CD: C-terminus domain B. Immunoblotting analysis of HEK293T cells. Cells were transfected with different clones of LRIG1 and stimulated with recombinant human EGF at indicated concentrations. C. Semi-quantification of the band in Figure 4B. D. Immunoblotting analysis of HEK293T cells. Cells were transfected with mock or LRIG1 vectors and stimulated with 10 ng/mL recombinant human EGF. The upper two blots are for proteins co-immunoprecipitated with EGFR antibody. The lower three blots are for whole lysates.

### Exon skipping event for exon 2 is detected in clinical cancer specimen and correlated with clinical outcome

To investigate whether LRIG1 variants with exon 2 skipping are expressed in patient tumors, we analyzed RNA-seq data from The Cancer Genome Atlas (TCGA) database. The exon 2 skipping event was evaluated by analyzing reads that connect exon 1 and exon 3 of LRIG1, as registered in TCGA SpliceSeq under the splice event ID: LRIG_ES_192431. In the SpliceSeq algorithm, Percent Spliced In (PSI) values are calculated for samples with at least eight alternative splicing-related reads.

Our analysis focused on six cancer types where more than 10% of samples were assigned PSI values: breast cancer (BRCA; 15%, N=181), esophageal squamous cell carcinoma (ESCA; 34.7%, N=67), glioblastoma (GBM; 23.1%, N=37), ovarian cancer (OV; 25.2%, N=106), prostate adenocarcinoma (PRAD; 11.1 %, N=61), and stomach adenocarcinoma (STAD; 34.3%, N=155). Among these cancer types, an average of approximately 3% of LRIG1 mRNA reads in the relevant region exhibited exon 2 skipping (LRIG_ES_192431), with certain cases exceeding 10% (Fig. 5A). Furthermore, cases with high expression levels of LRIG1 variants that lack exon 2 showed significantly poorer disease-specific survival in breast cancer and progression-free survival in ovarian cancer (Fig. 5B, C). These results suggest that the exon 2 skipping event in LRIG1 is not only detectable in various cancers but is also clinically relevant, potentially serving as a prognostic marker.

**Figure 5.**
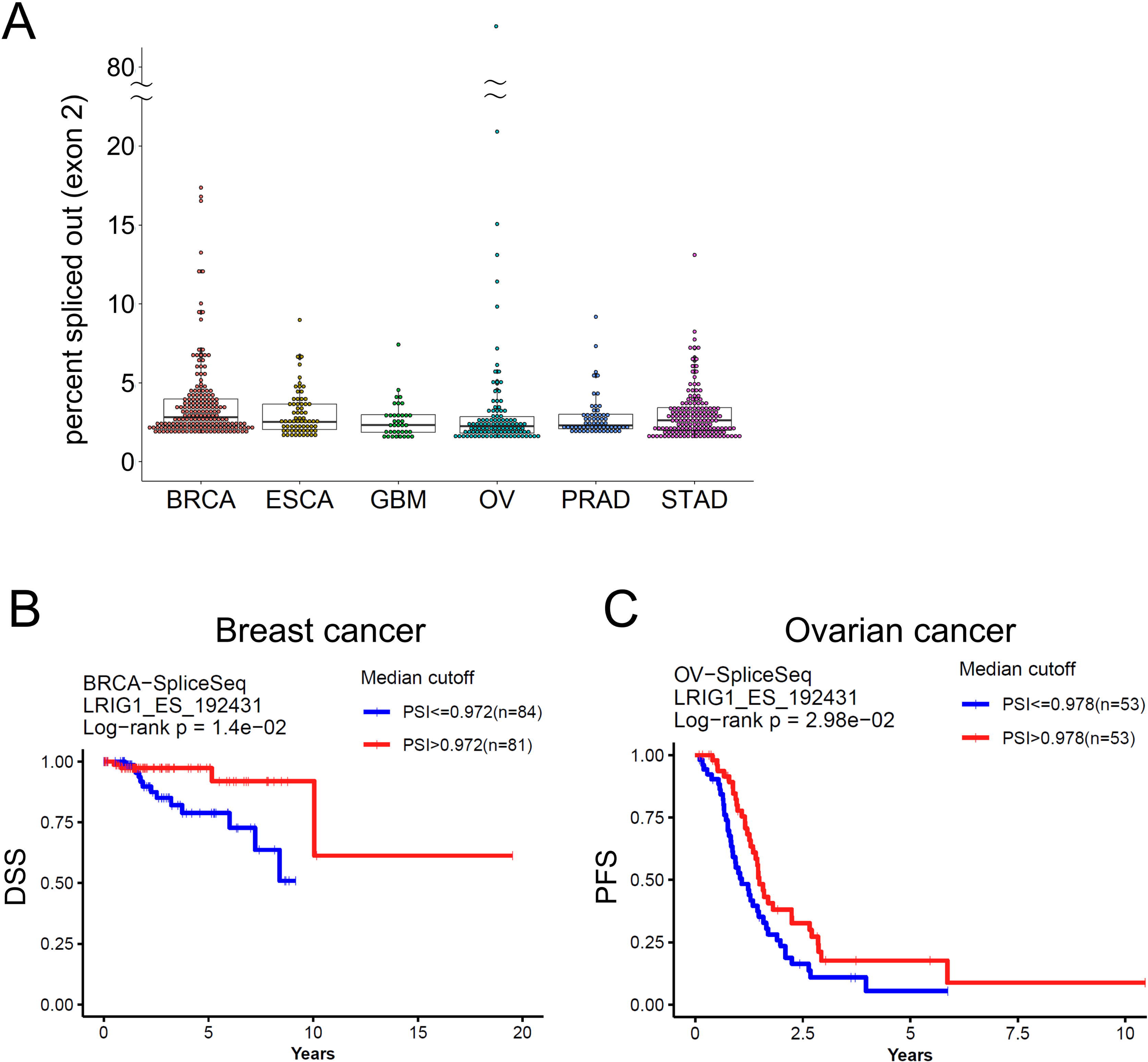
Exon skipping event for exon 2 is detected in clinical cancer specimens and correlated with clinical outcome. A. Graph showing the frequency of detection of LRIG1 exon 2 skip events in 6 cancers where PSI was assigned in more than 10% of all cases. Each dot indicates a case. BRCA: breast cancer, ESCA: esophageal squamous cell carcinoma, GBM: glioblastoma, OV: ovarian cancer, PRAD: prostate adenocarcinoma, STAD: stomach adenocarcinoma B. Kaplan-Meier curve of disease-specific survival (DSS) for patients with breast cancer stratified by frequencies of LRIG1 exon 2 skipping. A lower PSI (Percent Spliced In) value indicates a higher frequency of exon 2 skipping event (blue line) and vice versa for higher PSI (red line). Data registered in TCGA were analyzed by OncoSplice. C. Kaplan-Meier curves of progression-free survival (PFS) in patients with ovarian cancer stratified by frequencies of LRIG1 exon 2 skipping. Lower PSI indicates a higher frequency of exon 2 skipping event (blue line) and vice versa for higher PSI (red line). Data registered in TCGA were analyzed by OncoSplice.

## Discussion

In this study, we demonstrated that defective glycosylation of the membrane glycoprotein LRIG1 at its most N-terminal glycosylation site (N74 in humans) enhances EGFR binding and facilitates EGFR degradation. This finding highlights the previously unexplored role of specific glycosylation sites in modulating receptor-ligand interactions. Additionally, we identified that exon 2 skipping, a naturally occurring splicing event, generates a variant of LRIG1 that exhibits similar functional effects, further emphasizing the clinical relevance of alternative splicing in cancer biology.

The leucine-rich repeat (LRR) domain of LRIG1 is important for EGFR binding^7^, and the first five glycosylation sites from the N-terminus are located within this region. Among these, the N-terminal glycosylation site N74 is positioned on the concave surface, while N292 resides at the edge of the concave groove; the remaining three glycosylation sites are reported to be located on “one flank adjacent to the concave surface of LRIG1”^29^. Our data showed that the absence of glycosylation at N74 enhances LRIG1-EGFR binding, suggesting that the convex surface of the LRR domain directly interacts with EGFR while glycans at N74 may sterically prevent this interaction. Glycosylation, as a regulatory mechanism, positively modulates protein-protein interactions by influencing protein folding, stability, and binding specificity. For example, glycans can induce conformational changes or facilitate interactions with lectins, carbohydrate-binding proteins that recognize specific sugar moieties^11,14,30^. Conversely, some examples show glycosylation negatively regulates protein binding. For instance, glycosylation of the tetraspanin CD53 has been shown to inhibit interactions with its partner proteins CD45, CD20, and to a lesser extent, CD37^31^. Similarly, recent studies reported that the absence of glycosylation at specific sites on the ACE2 receptor enhances its binding to the SARS-CoV-2 spike protein^32,33^. These findings, along with ours, contribute to the growing body of evidence that glycosylation can act as a negative regulator of protein-protein interactions.

The introduction of gene mutations that eliminate glycosylation while minimizing the impact on the three-dimensional structure through N>Q substitutions is a widely employed method for investigating the functional significance of site-specific glycosylation in proteins. For example, in the case of the ACE2 receptor, this approach has been utilized to create artificial soluble ACE2 variants for therapeutic purposes, acting as decoys to capture SARS-CoV-2 viruses in the bloodstream. Mutants with specific N>Q substitutions have demonstrated enhanced viral capture efficacy^32^. However, glycosylation in cells is regulated globally, and no cellular mechanism has been identified that selectively eliminates glycans at specific sites in individual proteins. Additionally, no single nucleotide variants (SNVs) or cancer-specific mutations have been reported at the N74 position of LRIG1. In contrast, the results of this study revealed that a short exon containing the glycosylation site of LRIG1, exon 2, can be skipped while maintaining the reading frame. This splicing event results in the production of a LRIG1 variant with enhanced EGFR-binding capacity. To date, evidence linking the functional loss of glycosylation sites to exon skipping is limited. One such example involves the membrane receptor Neuropilin-1 (NRP1), which contains N-glycosylation sites at N150 and N261 in exons 4 and 5, respectively. Splice variants of NRP1 that lack exon 4 or exon 5 exhibit increased metastatic potential through enhanced interactions with Met and β1 integrin. This functional change was also reproduced in mutants with N150Q and N261Q substitutions, underscoring the role of glycosylation loss in regulating protein behavior through exon skipping^28^. These isolated examples illustrate the potential link between alternative splicing and glycosylation deficiency. Our findings in LRIG1 provide critical new insights into how alternative splicing contributes to the loss of N-glycosylation at key sites, resulting in functional alterations in protein-protein interactions.

Our database analysis revealed a correlation between LRIG1 exon 2 skipping and poor clinical outcomes in breast and ovarian cancers. While LRIG1 is known to suppress tumor growth through its role in EGFR degradation^34^, it also plays a critical role in maintaining quiescence in normal stem cells^4^. The downregulation of EGFR caused by exon 2 skipping in LRIG1 may induce quiescence in cancer stem cells, potentially leading to chemotherapy resistance or serving as the cell-of-origin for recurrence. Further exploration of the functional consequences of LRIG1 glycosylation status, particularly in the context of exon 2 skipping, may uncover its potential as a clinically relevant prognostic biomarker or therapeutic target in cancer.

## Acknowledgements

The authors thank Dr. Masahiro Inoue for providing colorectal cancer organoid C45. This study was supported by the Japan Society for the Promotion of Science Grant-in-Aid for Scientific Research (KAKENHI, 21K07942 (JK) and 22H02967 (EM)) and Osaka University Graduate School of Medicine Division of Health Science Management Grant.

